# Control of the force-bearing properties of microtubule-associated proteins via stalk/linker regions: insights from the NDC80 complex

**DOI:** 10.1101/2025.02.12.637486

**Authors:** Ilya B. Kovalenko, Ekaterina G. Kholina, Vladimir A. Fedorov, Egor M. Pozdnyakov, Philipp S. Orekhov, Nikita B. Gudimchuk

## Abstract

Many microtubule-associated proteins (MAPs) function under mechanical loads. Among them, motor proteins and passive couplers link microtubules with other cytoskeletal filaments, membranous structures and diverse scaffolds to enable cell shape changes, locomotion and other important processes. A key kinetochore complex, NDC80, transmits forces from microtubule disassembly to chromosome motion during cell division. Recently, this complex has been shown to detach from microtubules more easily when pulled toward the minus-end of the microtubule than when pulled in the plus-end direction. Here, we used coarse-grained molecular dynamics and Brownian dynamics simulations to explain the asymmetric effect of the directional load on the unbinding of the NDC80 complex from microtubules and then generalized our findings to other MAPs. We found that the lever arm created by the stiff stalk of NDC80 tilted toward the plus-end of the microtubule is critical for asymmetric unbinding of this complex, similar to that of dynein. In contrast, EB-proteins, the microtubule crosslinker PRC1, and kinesins are predicted to lack pronounced unbinding asymmetry, either due to their almost perpendicular anchorage to the microtubule wall or due to the high flexibility of their linker regions proximal to the microtubule-binding domains. Thus, our study highlights some of the design principles of MAPs, explaining how their distal parts can impart, modulate or eliminate the dependence of unbinding on the direction of external loads. This information deepens our understanding of the load-bearing properties and functions of diverse MAPs and may guide the design of synthetic protein systems with predefined mechanical characteristics.

## Introduction

Microtubule-associated proteins (MAPs) perform a plethora of physiological functions [1]. A considerable number of them have evolved to function under directional mechanical loads. For example, motor proteins, such as dynein and kinesin, are ATP-dependent force generators that carry loads along microtubules to their so-called minus and plus ends, respectively, enabling intracellular transport from the cell periphery to the center and vice versa [2]. Coupler proteins link microtubules to various intracellular structures, such as chromosomes, organelles, focal adhesion complexes and other scaffolds, enabling the segregation of genetic material, imparting mechanical stability to the cell, or mediating processes such as cell adhesion or migration [3]. Some notable examples of these MAPs include protein regulator of cytokinesis 1 (PRC1), a protein that crosslinks antiparallel bundles of microtubules to maintain the spindle midzone [4]; end-binding (EB) proteins that link various scaffolds to growing microtubule ends [5]; and spectraplakin family proteins that directly bundle microtubules with actin filaments [6]. Another instance is the NDC80 complex, a key kinetochore component that tethers chromosomes to microtubule tips during cell division to transduce mechanical force [7], [8], [9] (Fig. 1A). The Spc24 and Spc25 subunits of this heterotetrameric complex are associated with the inner kinetochore proteins. The globular domains of these two subunits are separated from the microtubule-binding interface via a ∼60-nm-long stalk that is mostly coiled-coil.

**Figure 1:**
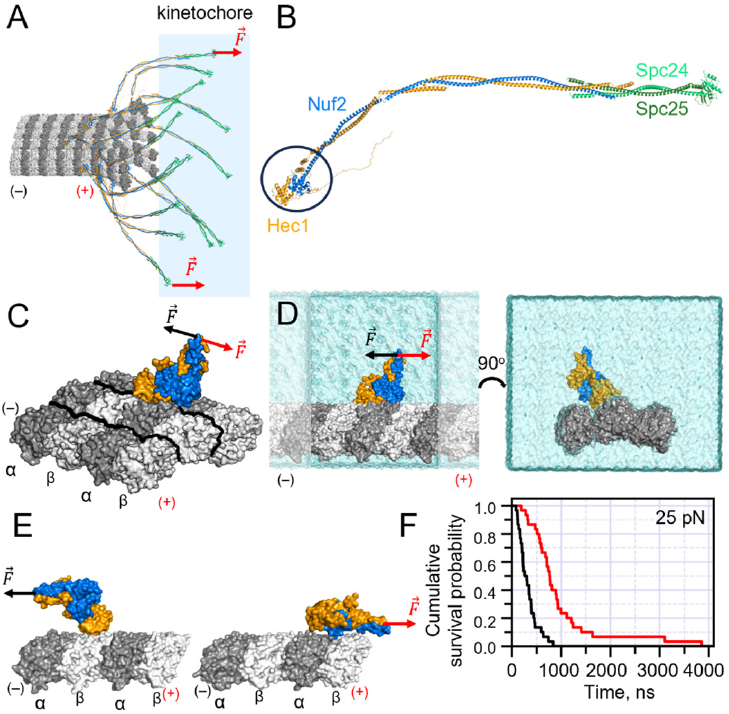
Modeling NDC80–microtubule interactions under plus- and minus-end-directed loads. **(A)** Schematic of chromosome-microtubule interactions mediated by NDC80 complexes. **(B)** AlphaFold2-predicted structure of the full-length NDC80 complex (50). **(C)** A CG model of a fragment of the NDC80 complex, containing Hec1 (orange) and Nuf2 (blue) subunits on a microtubule represented by a lattice of 3×4 tubulins. Black outlines mark boundaries between individual protofilaments. **(D)** Side and front views of the simulation box with periodic boundary conditions, filled with solvent. **(E)** Snapshots of the simulations of NDC80 unbinding from the microtubule when pulled with a plus- and minus-end-directed forces, applied to the tip of its coiled-coil stalk. **(F)** Kaplan-Meier plots, showing the cumulative survival probability for the NDC80 complex residence on the microtubule wall (based on *N* = 30 simulation runs for each load).

Microtubule binding is mediated by the globular calponin homology (CH) domain of the Hec1 subunit (Fig. 1B)[10], [7]. The fourth subunit, Nuf2, also affects microtubule binding, although the mechanism underlying its contribution is poorly understood [11].

Many critical insights into the functions and mechanisms of MAPs have come from single-molecule experiments. Optical trapping has been used to probe the motions of kinesins and dyneins with directional loads applied to the plus and minus ends of microtubules. The distinct force-velocity and force-detachment curves of kinesins and dyneins have been found to be tightly tuned for the optimal performance of these motors in their specific physiological contexts [12]. Interestingly, dyneins [13], [14], [15], [16] and kinesins possess an asymmetric unbinding rate dependence on the direction of a load, although the extent of asymmetry may vary for different kinesin family members [17], [18], [19], [20], [21]. In other words, both types of motor proteins are more easily detached when pulled in the direction of their natural motion along the microtubule, compared to pulling in the opposite direction. Analogous analyses of unbinding under directional loads for nonmotor proteins are scarce, although some proteins, such as EB1, PRC1, NuMa, and the Ska1 complex, have been shown to exhibit molecular friction when dragged along microtubules, with almost identical responses in both directions [22], [23]. Recently, using a novel three-bead assay and ultra-fast force spectroscopy, microtubule binding of the NDC80 complex has been shown to be sensitive to the direction of the applied load: its grip on the microtubule is an order of magnitude stronger when the complex is pulled toward the plus-end of the microtubule than when it is pulled to the minus-end [23], [24]. Similar behavior has also been observed for NDC80-containing artificially assembled minimal kinetochores and purified kinetochore particles [25]. This is intriguing, given that NDC80 binding to microtubules is mediated by the CH domain, which is structurally similar to the CH domain of EB proteins, but the latter do not display such unbinding asymmetry [22].

Here, we carried out extensive molecular dynamics (MD) simulations with the most recent version of Martini 3 general purpose coarse-grained (CG) force field refined for protein-protein interactions [26] and Brownian dynamics (BD) simulations to investigate the effects of directional load on protein unbinding from microtubules. We primarily focused on the NDC80 complex to determine the origin of its unusual behavior under force and then generalized the results to other MAPs. We found that the mechanism of asymmetry of the force response of the NDC80 complex is at least twofold. First, the force directed to the plus-end promotes contacts between the microtubule and the Nuf2 subunit, which presumably do not interact under zero load [27]. Second, we found that even a single Hec1 subunit alone displays significant asymmetry in microtubule residence time under load due to a lever-arm effect. This effect is created by a stiff coiled-coil stalk of the NDC80 complex, which is tilted to the plus-end at ∼45 degrees with respect to the microtubule axis. Finally, we showed that repositioning of this stalk orthogonally to the microtubule axis or making it flexible almost completely removed the dependence of microtubule residence time on the direction of the load. Our study highlights important design principles of MAPs finetuned by evolution, showing how the mechanical properties and geometry of their stalk/linker regions can impart, modulate, or eliminate the sensitivity of protein unbinding from its substrate to the direction of external load.

## Results

### Simulations recapitulate the asymmetric unbinding of the NDC80 complex from the microtubule

To investigate the molecular origin of the asymmetry of NDC80 unbinding from the microtubule under directional loads, we constructed a model of this complex on a microtubule wall. The model included a microtubule-proximal fragment of the NDC80 complex, containing the N-terminal regions of the Hec1 and the Nuf2 subunits, which included their CH-domains and short fragments of the alpha helices sufficient for dimerization (Fig. 1B). The CH-domains were positioned on a microtubule lattice, represented by 6 tubulin dimers arranged in 3 adjacent protofilaments (Fig. 1C), as informed by a cryoEM structure [27]. The protofilaments were effectively infinite across the chosen periodic boundary conditions (PBC). All unstructured regions of the proteins were excluded from consideration. The molecular system was consequently coarsegrained, minimized, and equilibrated in a rectangular simulation box with PBC applied as detailed in the Methods (Fig. S1, 1D). With this model, we examined the behavior of the NDC80 complex under a constant force of 25 pN applied at the tip of the NDC80 short stalk fragment in the plus- and minus-end directions along the microtubule (Fig. S2, Fig. 1E, Videos S1, S2). The simulations revealed an approximately 3-fold difference in the median NDC80 detachment time under directional loads (Fig. 1F). The unbinding rate was greater when the force was directed to the minus end of the microtubule, consistent with experimental data [24], [25].#

### Non-Hec1 domains contribute to tubulin–NDC80 interactions under load

We questioned whether the observed asymmetry of NDC80 behavior was a property of the microtubule-interacting Hec1 subunit or whether additional domains were also involved. Inspection of the simulation trajectories suggested that both the Nuf2 subunit and the stalk fragment transiently contacted the tubulins. However, the contacts between the stalk fragment and the tubulins appeared nonspecific and often involved the terminus of the truncated stalk, which is not be accessible in the full length NDC80 construct. Therefore, we decided to substitute the truncated stalk with a virtual stalk, represented by a number of stiff springs connecting a couple of virtual particles located about 6 nm above the origin of the stalk to the surface of the NDC80 globular domains (Fig. 2A). The equilibrium tilt angle and the length of the virtual stalk were set to mimic the orientation and length of the NDC80 stalk in the CryoEM structure [27]. This approach provided us with the following benefits, compared to using the explicit stalk in the simulation: (i) potential nonspecific interactions between tubulins and the stalk were eliminated, and (ii) we could easily change the length and orientation of the virtual stalk to study the effects of its geometry on NDC80 behavior under external forces.

**Figure 2:**
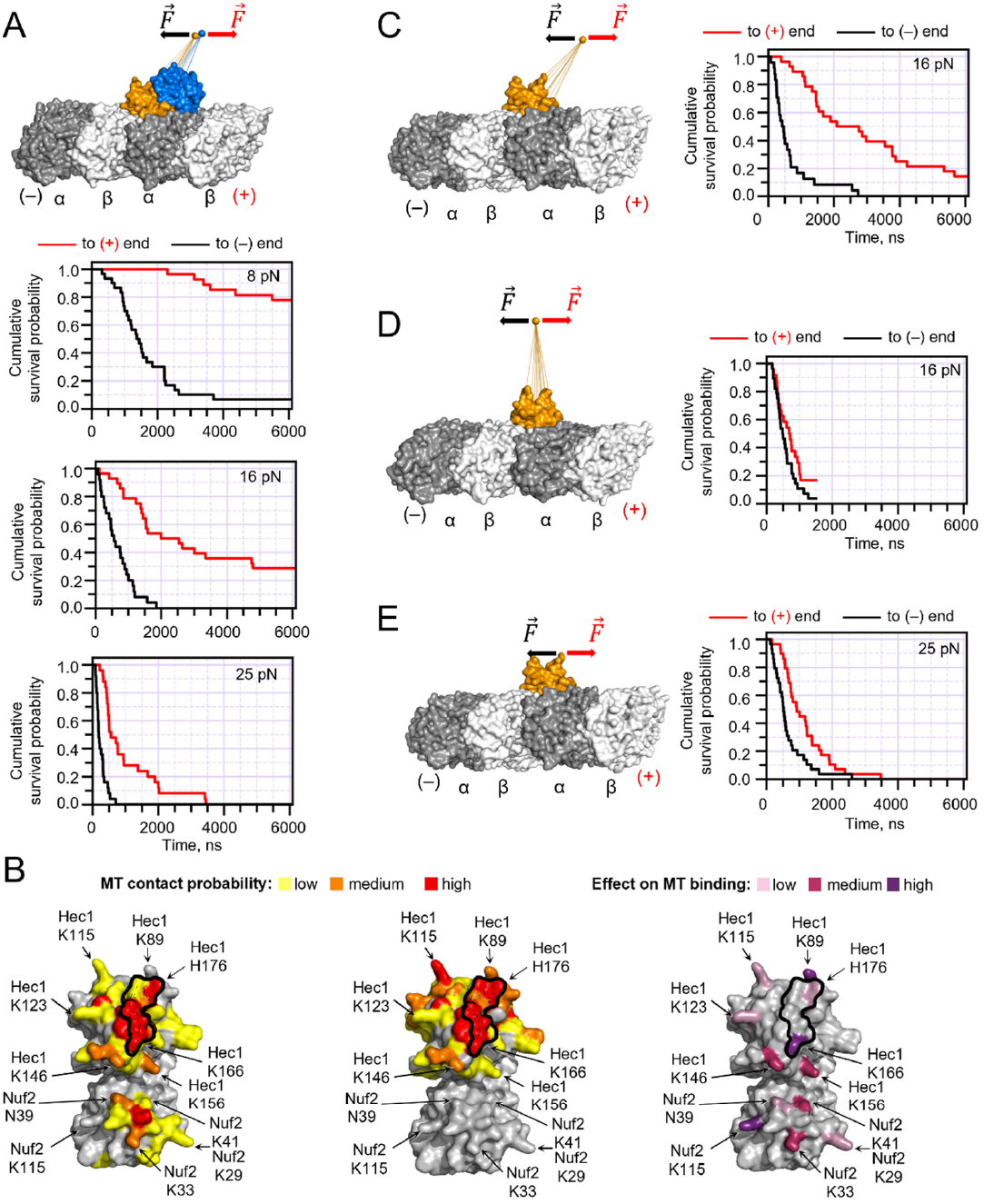
Role of the linker/stalk and Nuf2 subunit in NDC80 detachment from the microtubule. **(A)** Top: schematic of the simulations, in which the load is applied to the tip of the virtual stalk. The virtual stalk is represented by a set of springs connected to the CH-domains of the NDC80 complex. Bottom: Kaplan-Meier plots, showing the cumulative survival probability for the NDC80 complex residence on the microtubule wall in the CG MD simulations under directional loads: 8 pN, 16 pN, and 25 pN applied to the tip of the virtual stalk. Here and below the survival plots are based on at least *N =* 30 simulation runs for each condition. **(B)** Left: molecular surface of the Hec1/Nuf2-dimer colored on a gradient from yellow to red according to the probability of amino acid contact with tubulins under plus-end-directed load of 16 pN, viewed from the microtubule-binding interface. Middle: an analogous surface map, depicting contact probabilities under minus-end-directed load of the same magnitude. Right: a map of amino acids, whose mutations decrease the affinity of the NDC80 to microtubules, reproduced from (11). Black outline delineates the tubulin-binding interface in the CryoEM structure (27). **(C-E)** Simulation schematics and Kaplan-Meier plots, showing the cumulative survival probability for the residence of a single Hec1 subunit on the microtubule wall in the CG MD simulations: **(C)** the load magnitude is 16 pN and the virtual stalk is in the naturally tilted position, **(D)** the load is 16 pN and the virtual stalk is orthogonal to the microtubule, **(E)** the load is 25 pN and the load is applied to the origin of the stalk.

The simulations revealed that load application via a ∼6-nm-long virtual stalk reproduced the unbinding asymmetry (Fig. S2B, Fig. 2A) observed in the simulations with the explicit stalk (Fig. 1E,F) and in accordance with the experimental data [24], [25]. We found that force-driven tilting of the NDC80 complex promoted a handful of additional contacts between the Hec1 subunit and the microtubule, which were not observed in the starting CryoEM structure (Fig. 2B). Moreover, plus-end directed, but not minus-end-directed loads facilitated contacts between the Nuf2 subunit and the microtubule. The additional contacts were consistent with the previously reported involvement of positively charged amino acids on both Hec1 and Nuf2 in microtubule binding [11]. This offers a possible explanation for the involvement of Nuf2 in interaction with tubulins, despite the lack of such contact in the CryoEM structure [27].

### A tilted stalk is sufficient to drive the asymmetric unbinding of a single Hec1 subunit

To clarify whether the Nuf2 subunit was the only driver of the observed unbinding asymmetry, we modified our model by excluding Nuf2 from it and attaching the virtual stalk only to the Hec1 subunit (Fig. S3A, Fig. 2C). In this setup, Hec1 alone displayed a distinct asymmetry of response to load (Fig. 2C). In the absence of the Nuf2 subunit, the tail of the distribution of detachment times under the plus-end-directed loads became lower while the median detachment times in both directions remained the same. Most importantly, repositioning of the virtual stalk orthogonal to the microtubule axis almost completely removed the unbinding asymmetry (Fig. S3B, Fig. 2D). This suggested that unequal unbinding times for plus- and minus-end-directed loads resulted mainly from the tilted equilibrium orientation of the stalk, creating a lever arm effect. Consistently, when the forces were applied directly to the point of origin of the stalk, the lever-arm effect almost disappeared, leading to only modest unbinding asymmetry (Fig. S3C, Fig. 2E).

### The orientation and flexural rigidity of the stalk govern the behavior of MAPs under load

Appreciating the role of the lever arm effect on NDC80’s unbinding from the microtubule prompted us to investigate the impact of stalk properties on MAP–microtubule interactions in the context of directional loads more broadly. We queried the Protein Data Bank for MAP microtubule structures with elongated stalk domains. In addition to the NDC80 complex, our analysis revealed two more prominent examples of MAPs having this feature: the microtubule crosslinker PRC1 [28] and the minus-end-directed motor dynein [29] (Fig. 3A). The majority of other MAPs appear to bind microtubules without creating any significant lever arms, either by lacking stalk regions protruding away from microtubules or by having linkers with substantial flexibility. For example, EB proteins have similar CH domains, like Hec1/Nuf2 of NDC80, but their CH domains extend into 58-residue-long unstructured linkers [30]. Due to their flexibility and length, these linkers should align the successive coiled-coil fragment of an EB protein along the direction of applied force, regardless of the attachment point of the linker on the CH domain. In other words, the linkers are expected to completely eliminate the ‘lever-arm effect’ (Fig. 3A). Analogously, the motor domains of kinesins are connected to the stalk with rather flexible neck linker regions and are also expected to have considerable flexibility around the onset of the stalk [31] (Fig. 3A).

**Figure 3:**
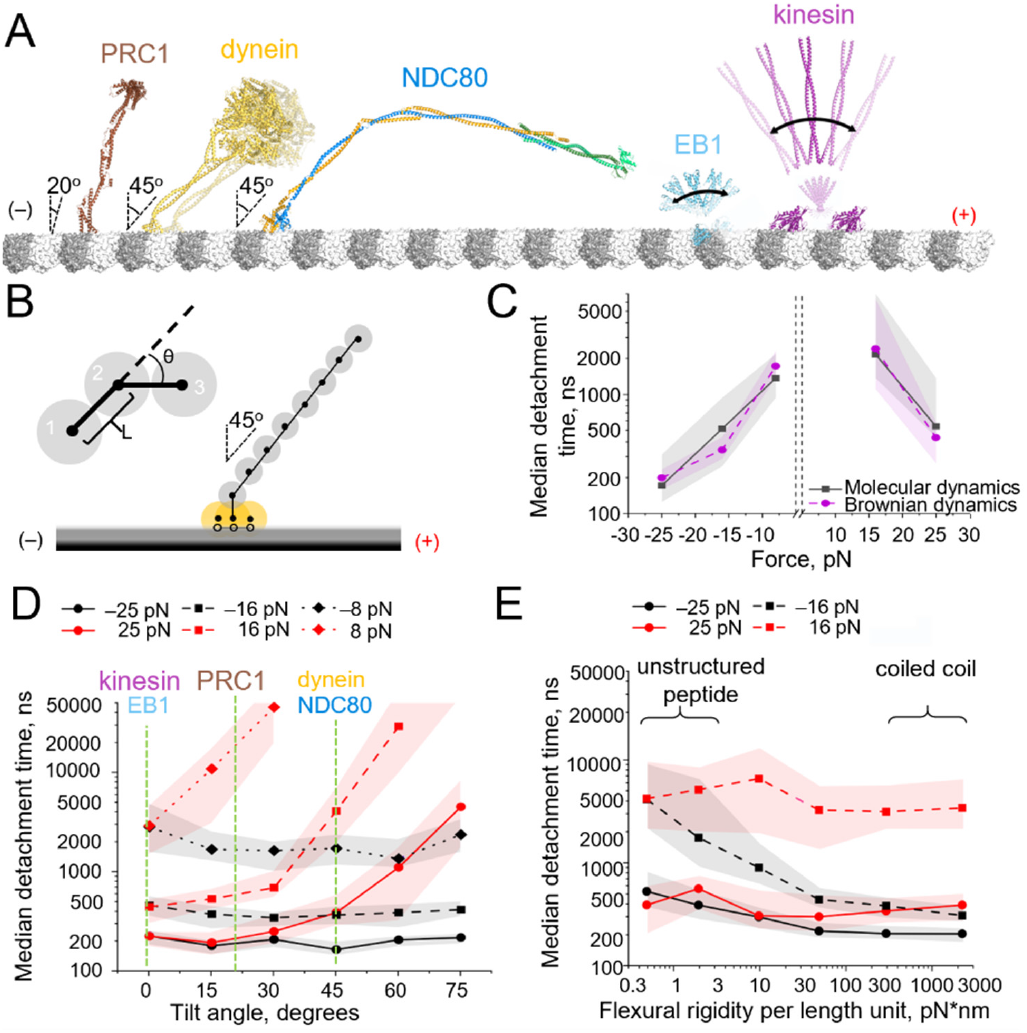
BD model of MAP-microtubule interactions. **(A)** Structures of five MAPs bound to the microtubule. For PRC1 the model shown is an overlay of PDB structures: 4L6Y [32] and 5KMG [28]; for dynein: an overlay of 4RH7[33] И 6RZB[34]; for NDC80: an overlay of PDB 3Iz0 [27] and AlphaFold2-prediction [35]; for EB1: 3JAK[36] and 1YIB[37]; for kinesin and motor domains: from PDB 3J6E[38], and the coiled coils 1 and 2 based on a recent study [39]. **(B)** Schematics of the BD model of MAP bound to the microtubule. Inset shows the bending potential, maintaining alignment of the beads, representing the stalk. **(C)** Parametrization of the BD model for NDC80 complex based on CG MD simulations. **(D)** Dependence of the NDC80 detachment times on the angle of the stalk relative to the microtubule in BD simulations. Median and interquartile range are shown based on *N* = 30 simulation runs for each point. **(E)** Dependence of the NDC80 detachment times on the flexural rigidity of the stalk in the BD simulations. Median and interquartile range are shown based on *N* = 30 simulation runs for each point.

To explore the impact of the stiffness and geometry of the stalk on the response of different MAPs to directional loads, we designed a simple Brownian dynamics (BD) model of a MAP attached to the microtubule wall (Fig. 3B). In this model, the microtubule interacting domain of the MAP was represented by three spheres (Fig. 3B, shown in orange), which were connected to a chain of smaller spheres, effectively modelling the stalk (Fig. 3B, gray). The energy potentials between the spheres in the chain maintained the curvature of the chain and the spacing between the spheres. The motions of the spheres were described by the overdamped Langevin equations [40]. We first parameterized the BD model to have the flexibility of the stalk, as reported for coiled coils [41], and the dimensions and orientation of the stalk, to mimic the cryo-EM structure of the NDC80 complex [27]. In this setup, the BD model successfully recapitulated the asymmetry of NDC80 unbinding times as a function of plus- and minus-end-directed loads. By tuning the free parameters of the BD model (see Methods), it was possible to achieve a behavior, corresponding to that of the NDC80 complex in our CG MD simulations (Fig. 3C, Video S3). We used the calibrated BD model to explore the impact of the stalk properties, such as the tilt angle relative to the microtubule axis, flexural rigidity and length, on detachment under directional loads. As expected, the extent of asymmetry in the unbinding rate increased as the orientation of the stalk deviated from orthogonal to tilted along the microtubule (Fig. 3D). Assuming a typical flexural rigidity of a coiled coil [41], the unbinding asymmetry was predicted to be modest when the stalk was slightly tilted, e.g., as in the case of PRC1 (∼20°) (Fig. 3D). The asymmetry was pronounced for stalks tilted such as those of NDC80 and dynein. When a considerable tilt was present, the flexural rigidity of the stalk was critical for maintaining asymmetry (Fig. 3E). For example, with flexible linker regions (as expected for EB proteins and kinesins), no asymmetry was predicted (Video S4), but with stiffer linkers, such as in NDC80 and dynein, the asymmetry was pronounced. The impact of the length of the stalk on the detachment time was more complicated because the length increased the leverage but also decreased the effective bending stiffness of the stalk and augmented the viscous drag of the protein, making its detachment slower. As a result, longer stalk regions increased the unbinding times in both directions (Fig. 4). For example, when we considered the full geometry of the nearly 60-nm-long NDC80 complex, it was clear that the viscous drag dominated the time of its unbinding from the equilibrium tilted state (Video S5).

**Figure 4:**
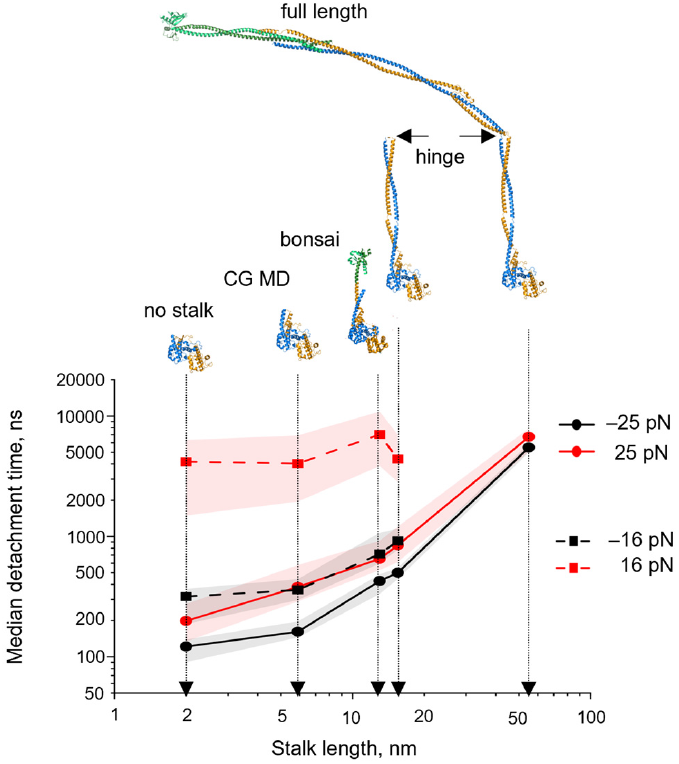
Dependence of the unbinding kinetics on the length of the stalk in BD simulations. Each data point shows a median and an interquartile range based on *N* = 30 simulation runs. The upper row of images shows fragments of the NDC80 complex corresponding to a given stalk length.

## Discussion

The extensive CG MD and BD simulations presented here collectively cover over 1 ms and 10 ms of simulated times, respectively (equivalent to almost 1 M cpu*hours of computation time). This timescale is sufficient to estimate NDC80 detachment rates under close-to-physiological loads, offering several mechanical insights into the interactions between this kinetochore complex and the microtubules.

First, we identified the possibility of direct involvement of the Nuf2 subunit in interactions with microtubules under a plus-end-directed force. Despite the limitations of our CG MD approach restricting flexibility of the spatial structure of proteins, the predicted major binding region of the Nuf2 subunit to tubulin aligns well with the previous *in vitro* analysis [11], which mapped mutations in the Nuf2 subunit, that led to decreased microtubule-binding affinities of Ndc80. Together, these data indicate that Nuf2 may be directly involved in the interactions with the microtubule under external load. It is also possible that such contacts are transiently present in equilibrium but are not observed in the experimental CryoEM structures [38], [42], perhaps due to their dynamic nature in the absence of external load. Thus, we speculate that the application of a plus-end-directed force, which tilts the Nuf2 subunit toward the microtubule, could enhance overall NDC80 complex binding at a certain range of forces. This could explain the previously documented but poorly understood catch-bond behavior of purified kinetochore particles[43]. Unfortunately, we were unable to explore the range of small forces where catch-bond behavior is typically observed due to computational limitations. Investigating this regime remains an exciting prospect for future work.

Second, our simulations emphasize the role of the stalk/linker regions in modulating the response to the load of NDC80 and other MAPs. Two factors, the equilibrium tilt of the stalk along the microtubule, and the flexural rigidity of this region, are found to be critical for causing an asymmetric response to load. A close-to-orthogonal orientation of the stiff stalk relative to the microtubule axis, such as in PRC1, or a highly flexible linker/hinge region, such as in kinesins or EB proteins, are not predicted to confer any unbinding asymmetry. It should be noted, however, that the asymmetric response to load may also be caused by other factors. As one example, altering the neck linker orientation relative to the motor domain of kinesin-1 by the plus-or minus-end-directed loads was reported to determine the nucleotide binding, thereby controlling the affinity of the motor domain to the microtubule track [19].

Another conceivable determinant of asymmetric unbinding of a MAP from the microtubule could be a nonuniform distribution of the strength of microtubule contacts within the interaction interface. This factor may be responsible for residual unbinding asymmetry in our CG MD simulations with NDC80 in the absence of the virtual stalk (Fig. 2E), while it is negligible for unbinding of the CH-domain of EB3 (Fig. S3D-F). A modulation of the unbinding asymmetry by interface amino acids has been recently described for dynein, where a single mutation at the microtubule binding interface was sufficient to remove the unbinding asymmetry of dynein [44]. Consistently, we recapitulated this effect in our BD model, in which the three-point microtubule interaction potential was redistributed to create a deeper potential well on the side (Fig. 5, Video S6). In that case, we observed an unbinding asymmetry even when the stalk was oriented orthogonally at equilibrium.

**Figure 5:**
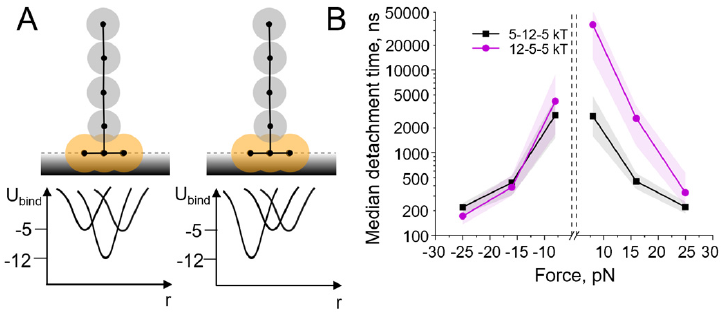
Role of the distribution of the strength of binding interactions along the tubulin binding interface. **(A)** Schematics of the starting configuration of the BD simulation. Left: the interaction points are positioned symmetrically, with the strongest potential well, 12 k_B_T, in the middle, and weaker potential wells, 5 k_B_T, on the sides. Right: the interaction points are re-distributed so that the strongest interaction point is at the left side (shifted to the minus end of the microtubule). **(B)** Detachment times in the symmetric and asymmetric cases under plus- and minus-end-directed loads. Each data point shows a median and an interquartile range based on *N* = 30 simulation runs.

What are the physiological implications of the asymmetric unbinding feature of MAPs, which possess a tilted and rigid stalk? NDC80 and dynein are important mitotic proteins that perform a number of critical functions throughout cell division (Fig. 6A).

**Figure 6:**
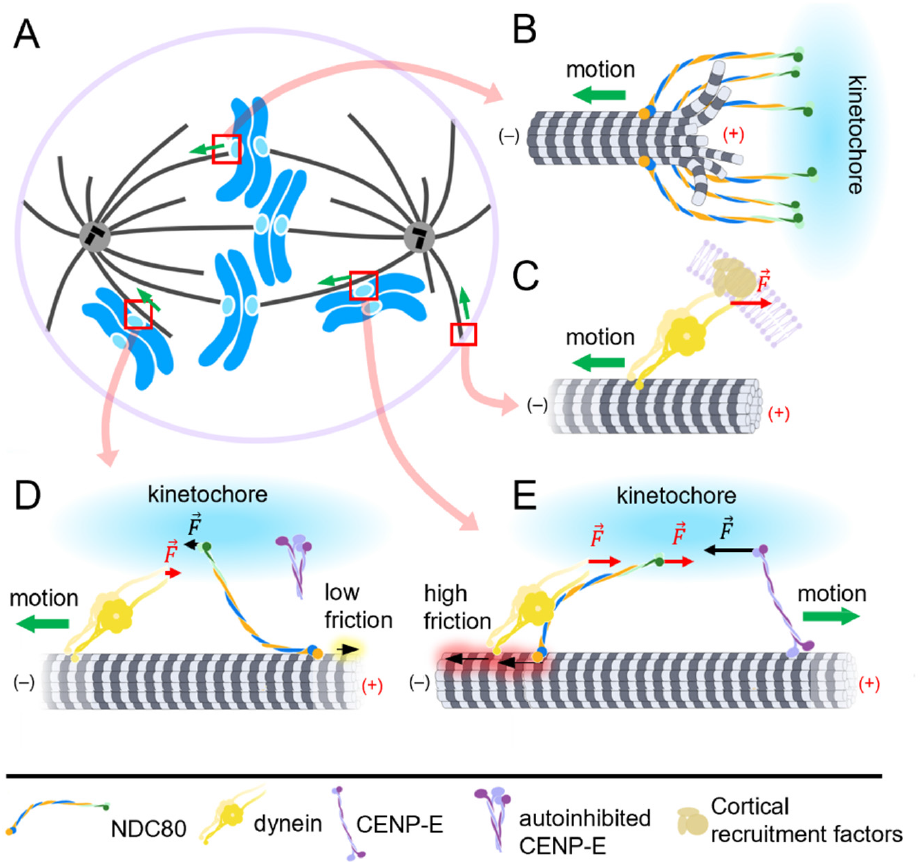
Possible roles for asymmetric unbinding of NDC80 and dynein in mitosis. **(A)** Schematics of a dividing cell. The directions of chromosome or microtubule motions are shown with green arrows. **(B)** Microtubule end attached to the kinetochore via NDC80 complexes, maintaining a strong grip when pulled with plus-end-directed force from the sister kinetochore. **(C)** Cortical dynein maintaining a strong grip on the astral microtubule plus-end under plus-end-directed load. **(D)** A laterally attached kinetochore is moved by dynein in the minus-end direction (toward the spindle pole). In this case, NDC80 displays a weak grip on the microtubule, leading to low molecular friction; kinesin-7 (CENP-E) is autoinhibited. **(E)** A laterally attached kinetochore is moved by CENP-E in the plus-end direction along the spindle microtubule bundle. NDC80 displays a stronger grip on the microtubule, leading to high molecular friction. The strength of this attachment is likely downregulated by its phosphorylation at the pole. Dynein is also in a ‘high friction mode’; albeit we hypothesize its resistance may be similarly downregulated through a yet unknown mechanism.

NDC80 serves as the primary coupler of microtubule tips to kinetochores, so its ability to strongly grip the plus-ends and persistently maintain attachment to microtubules under plus end-directed loads is of clear physiological value (Fig. 6B). Similarly, kinetochore-associated dyneins may also contribute to the grip of the microtubule plus-ends, although the extent and importance of this contribution is debated [45]. More established is the role of cortical dyneins in anchoring the tips of astral microtubules to the cell membrane [46]. This is essential for positioning mitotic and meiotic spindles [47]. The geometry of the dynein stalk, tilted toward the plus end of the microtubule, is beneficial for a stronger grip in this setup (Fig. 6C). The ability of NDC80 and dynein to detach from microtubules asymmetrically, i.e. more easily, when pulled in the minus end direction rather than the plus-end direction may also have important physiological implications. First, this feature is thought to contribute into directionality of dynein’s stepping: the trailing foot of dynein experiences a minus-end-directed load during stepping, so it detaches faster than the leading foot, which experiences a plus-end-directed load [13], [14], [15], [16]. Second, we propose that the asymmetric unbinding of NDC80 under load may enable direction-selective resistance to orchestrate chromosome motions in prometaphase following nuclear envelope breakdown. Although some chromosomes rapidly become end-on attached to microtubules, the initial capture of peripherally located chromosomes frequently relies on lateral contacts between the microtubule and kinetochores [48]. Despite the simultaneous presence of NDC80 complexes and two oppositely directed motors, dynein and kinesin-7 (CENP-E), at the kinetochore, these proteins do not appear to engage in an unproductive tug of war, but first allow minus-end-directed motions to bring the chromosomes to the spindle pole and then deliver them to the cell equator via plus-end-directed motions along preexisting spindle microtubules [49]. This sequence of events has been shown to be important for correct chromosome segregation [49], [50]. How is this regulation achieved? CENP-E has recently been suggested to reside at the kinetochore in a self-inhibited state at the onset of prometaphase [51]. Thus, dynein dominates the initial chromosome transport, moving the chromosomes in the minus-end direction toward the spindle pole. Importantly, NDC80 does not impede dynein-mediated transport due to the rapid unbinding of NDC80 from the microtubule when its stalk is tilted toward the minus-end (Fig. 6D). Upon reaching the spindle pole, CENP-E and NDC80 are phosphorylated by the centrosome-localized Aurora A kinase, which presumably alters the balance of forces at the kinetochore, leading to the dominance of CENP-E-driven chromosome transport (Fig. 6E). Specifically, phosphorylation has been reported to activate CENP-E by relieving its autoinhibition [52], [51]. Based on in vitro evidence, NDC80 has been proposed to act as a brake for CENP-E-mediated chromosome transport [53]; however, partial phosphorylation of NDC80 at the pole likely helps to decrease frictional drag [54], [55]. Thus, the mechanical properties of NDC80 are fine-tuned to allow proper direction and order of chromosome motion for successful delivery to the metaphase plate.

Overall, our study highlights the role of “passive” regions, such as linkers and stalks, in the dynamical behavior of protein complexes, which have been largely neglected in the literature, while the major focus has been on more “active” factors, such as microtubule affinity or activity of motor domains. Understanding the mechanics of these regions adds to our understanding of MAP-microtubule interactions and illuminates the principles for rational design of artificial proteins with asymmetric and symmetric responses to load.

## Acknowledgements

We are grateful to Fazly Ataullakhanov for discussions and important ideas, J.R.McIntosh for reading the paper, and Julia Borzunova for technical assistance. We thank Michael A. Cianfrocco for providing a model structure of autoinhibited kinesin-1. This work was supported by the Russian Science Foundation grant # 23-74-00007 to I.B.K. Simulations were partially carried out using the equipment of the shared research facilities of HPC computing resources at Lomonosov Moscow State University. P.S.O. is a member of an innovative drug development team based on structural biology and bioinformatics at Shenzhen MSU-BIT University #2022KCXTD034. This paper was typeset with the bioRxiv word template by @Chrelli: www.github.com/chrelli/bioRxiv-word-template

## Author contributions

I.B.K., E.G.K., V.A.F. performed and analyzed the coarse-grained molecular dynamics simulations. E.M.P. performed and analyzed the Brownian dynamics simulations. P.S.O. supervised and analyzed the coarse-grained simulations, and edited the manuscript. N.B.G. designed the study, analyzed the data, and wrote the manuscript.

## Materials and Methods

### CG MD simulations

We constructed five molecular systems representing different fragments of a single copy of the NDC80 complex, using the PDB structure 3IZ0 [27] and a short segment of the microtubule wall from the PDB structure 5JCO [56], which consisted of three protofilaments, each composed of two tubulin dimers. The systems included: (1) a fragment of the NDC80 complex with a ∼6 nm-long coiled-coil stalk; (2) globular domains of Hec1/Nuf2 with a virtual stalk, mimicking a ∼6 nm-long coiled-coil; (3) globular part of only the Hec1 subunit of the NDC80 complex with a virtual stalk mimicking a ∼6 nm-long coiled-coil stalk; (4) the same system with an orthogonally positioned virtual stalk; and (5) the same system without any stalk. Additionally, we constructed a molecular model of the EB3 calponin homology domain based on the PDB structure 7SJ9 [57], which is associated with a short segment of the microtubule wall based on the PDB structure 5JCO [56]. Each of the molecular systems were converted to CG representation using the martinize2.py workflow module of the MARTINI 3 force field [26] (https://github.com/marrink-lab/ver-mouth-martinize) considering the secondary structure DSSP assignment [58]. We used the ElNeDyn elastic network approach [59] with a default force constant of 500 kJ/(mol·nm^2^) to maintain the stability of the secondary structure of the modeled proteins. The pH of the systems was considered neutral. All the simulations were run in the presence of regular MARTINI water, neutralized, and ionized with Na^+^ and Cl^−^ ions at a concentration of 150 mM. The systems were equilibrated for 200 ns. We utilized the reaction-field approach for long-range electrostatics with a cutoff of 1.1 nm, and the dielectric constant beyond the cutoff was set to infinity as recommended in [60], [61]. All simulations were conducted in a NVT ensemble due to the presence of the positional restraints using the V-rescale thermostat at 310 K with a coupling constant of 1.0 ps. The integration time step was 20 fs. A constant force of 9 pN, 16 pN, or 25 pN was applied to the NDC80 at various application points and in the opposite directions along the microtubule axis. In experiments with EB3 a constant force of 100 pN was applied to the origin of the flexible linker. CG MD simulations were performed using GROMACS version 2022.4, which allows parallel computing on hybrid architectures [62].

The simulation trajectories were processed using Python 3 scripts to determine how the number of remaining native contacts and new contacts between NDC80 and tubulins changed over time in each simulation. We used the native contacts analysis method implemented in the MDAnalysis Python library [63]. Native contacts of Hec1 with microtubule tubulin were defined as the proximity within a 5 Å distance threshold, based on the starting configuration. We applied the *soft_cut* option with a softness parameter of 2.0 and a reference distance tolerance of 3.5. A detachment event was defined as the rupture of more than 80% of the native contacts. The cumulative survival probability for the residence of the NDC80 complex on the microtubule wall was represented by Kaplan-Meier plots. New contacts were calculated analogously, based on subsequent simulation frames. Probabilities of contacts between Hec1 or Nuf2 and tubulins throughout the MD trajectory were calculated as the average fraction of contacts over the entire simulation time.

### BD simulations

Our BD model of a MAP consisted of three spheres representing the tubulin-interacting interface of the microtubule-binding domain (MBD), and a chain of smaller spheres, the first one representing the back of the MBD and the rest representing the flexible stalk.

Each of the three tubulin-interacting spheres could bind to a corresponding site located on the line representing the microtubule surface via a Gaussian-shaped potential:

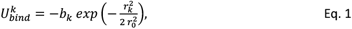

where *k* = 1, 2, 3 describes the position of the binding site for the NDC80, ordered from the minus end to the plus end; *b*_*k*_ is the corresponding depth of the *k*-th interaction potential; *r*_*k*_ is the distance between the center of the *k*-th of the MBD sphere and its corresponding binding site, and *r*_*O*_ is the width of the potential well.

The sphere, representing the back of the MBD, was flexibly linked to the spheres representing the tubulin-interacting interface and to the first element in the chain of the spheres, representing the stalk. Linear and angular deformations of the stalk were controlled by quadratic energy potentials. Linear deformations were determined as the change in distance between the centers of two neighboring spheres while angular deformations were elicited from the angle between the lines connecting the centers of adjacent pairs of spheres within the stalk:

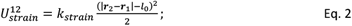

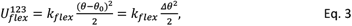

where ***r***_*1*,*2*_ are the coordinates of the neighboring spheres of the stalk, *l*_*O*_ is the distance between the adjacent sphere centers when the stalk is not deformed, *θ* is the angle between the two lines connecting the three spheres, and *θ*_*O*_ is the value of this angle when the stalk is not deformed, *Δθ* is the angular deformation of the stalk.

A constant force was applied to the center of the distal bead in the plus-or minus-end direction along the microtubule, evoking the following energy potential:

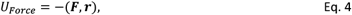

where ***F*** is the force vector and ***r*** is the radius vector of the distal bead of the stalk.

Any part of the MAP was expelled from the space below the line representing the microtubule surface by a quadratic repulsion potential:

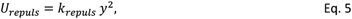

where *y* is the depth of the sphere center penetration into the hemispace below the microtubule.

Thus, the total energy of the system, *U*, is defined as the sum of the following terms:

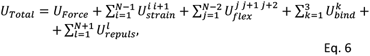

where *N* is the total number of spheres that move independently of one another.

The motions of the spheres were simulated by solving an overdamped Langevin equation [40]:

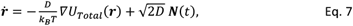

where 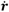 is the velocity of the sphere, *D* is the diffusion coefficient of the sphere, *k*_*B*_ is the Boltzmann constant, *T* is the temperature of the system, *∇U*_*Total*_(***r***) is the system’s potential energy gradient with respect to the coordinates of the sphere whose movement is being described, and ***N****(t)* is a white noise term.

The diffusion coefficient for each sphere was calculated as:

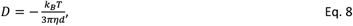

where *η* is the viscosity coefficient, *d* is the diameter of the sphere.

The simulations were run in Matlab R2022a. The MAP was considered detached when the distance between all of the pairwise interaction points on the MBD and the microtubule exceeded a cutoff of 1.6 nm (approximately two Debye lengths). Computer code is available upon request.

### BD model parameterization

A summary of default BD model parameters is provided in Table S1. The default parameters of a MAP were chosen to mimic the geometry and mechanics of the NDC80 fragment having an approximately 6-nm-long stalk, as in the PDB structure 3IZ0 [27]. Specifically, the diameters of the large spheres, representing the MBD of the NDC80, and the distance between the centers of the first and third spheres were set to *d*_*MBD*_ = 2.3 nm, and *h*_*MBD*_ *=* 1.8 nm, respectively. The back of the MBD and the ∼6 nm stalk were represented by four spheres with a diameter of 2 nm, corresponding to the width of a coiled coil. The stalk tilt angle was set to 45°.

The axial and flexural stiffness of the stalk were determined based on reported parameters of the coiled coil (see the section below).

For generality, the three tubulin-MBD interactions were described by a set of three independent parameters {b_1,_ b_2,_ b_3_}. By default, we assumed a symmetric distribution of the depths of the binding potentials: *b*_*1*_ = *b*_*3*_ < *b*_*2*_. The parameters of the tubulin-MBD interactions, {*b*_*k*_}, were calibrated to reproduce the ratios of MAP detachment times under the plus and minus-end-directed forces in CG MD simulations. The binding potentials parameters were then fixed and the viscosity parameter was varied to fit the absolute values of the MAP detachment times in the CG MD simulations. A good correspondence between the BD and the CG MD models was achieved at viscosity *η* = 4.8 mPa·s, which is ∼7-fold higher than the dynamic viscosity of water. This difference likely reflects an effective increase in viscosity in the CG MD model due to (i) peculiarities of the water model in Martini 3, which is bulkier as it substitutes 4 water molecules, resulting in a self-diffusion rate slower by a factor of ∼5 [64] and presumably higher viscosity of this water model, and (ii) the contribution of hydrodynamic effects, which were not accounted for in the BD model.

To explore the load-dependence of the MAP detachment time as a function of the mechanical parameters of the stalk, we varied the tilt angle, flexural stiffness, and length of the stalk (with the number of beads in the stalk changed accordingly), while keeping the other parameters at default values, described in Table S1. To explore the load-dependence of the MAP detachment time as a function of the distribution of the depths of the binding potentials, we swapped the binding potentials of the plus-end proximal and the central interaction points: {*b*_*1*_, *b*_*2*_, *b*_*3*_} = {5, 5, 12} k_B_T and set the tilt angle of the stalk to zero (orthogonal to the microtubule axis).

To estimate parameter *k*_*flex*_, which relates the energy of bending deformation of the stalk unit with its angular deformation (Eq. 3), the same energy was expressed as the function of the deflection magnitude, *w*, and the corresponding bending stiffness *κ* :

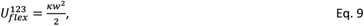

where *w* is the deflection distance of the 2nd sphere from the line, connecting the centers of the first and third spheres. Assuming the spheres are parts of a simply supported beam with a central load, we can re-write Eq.9 as:

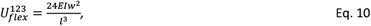

Here *E* is the Young’s modulus of the beam, and *I* is the second moment of area, *l* is the length of the beam unit. Assuming a small angular deformation, the deflection *w* can be expressed as follows:

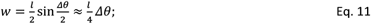

Combining Eq. 10 and Eq .11, we have:

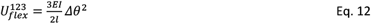

Comparing Eq. 12 and Eq. 3 yields an expression that links the bending rigidity, *EI*, per unit length *l* of the beam, with the harmonic stiffness, *k*_*flex*_:

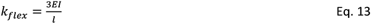

Given the relationship between the bending rigidity and the persistence length *P*:

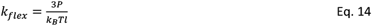

The axial stiffness of a unit length of the stalk can be expressed as:

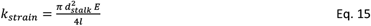

Formulas 14 and 15 allow estimating the mechanical parameters of the BD model based on published quantifications of the properties of the coiled coil, such as the persistence length *P* = 130-500 nm [65], [66], [67], [68], and Young’s modulus for stretching *E* = 2.7·10^9^ - 3.3·10^11^ J·m^-3^ [41], [69].

