## Supporting Information for "Control of the force-bearing properties of microtubule-associated proteins via stalk/linker regions: insights from the NDC80 complex"

**Table S1: BD model parameters.**
Default values are given. Several parameters were varied to explore their impact on model behavior

| Parameter | Description | Value | Source |
|---|---|---|---|
| $h_{MBD}$ | Height of the MBD | 1.8 nm | Based on dimensions of the MBD of NDC80 (PDB 3iz0) [1] |
| $d_{MBD}$ | Diameters of the spheres of the MBD | 2.3 nm | |
| $d_s$ | Diameter of a sphere in the chain representing a stalk | 2 nm | Equals the diameter of a coiled coil [2], [3] |
| $\{b_1, b_2, b_3\}$ | Depths of the energy potentials of the three pairs of interaction centers on the MBD and tubulins | {5, 12, 5 } $k_BT$ | Default values calibrated to reproduce CG MD simulations |
| $r_0$ | Width of the MBD-tubulin potential well | 0.8 nm | Estimated to approximately equal the Debye length in the cell [4] |
| $\theta_0$ | Equilibrium stalk curvature | at the joint between the MBD and the stalk: 45°<br><br>elsewhere: 0° | Default value for NDC80, based on the PDB structure 3iz0 [1] |
| $k_{flex}^{stalk}$ | Flexural stiffness of the stalk and MBD per unit length | 2300 pN·nm | Default value for a coiled coil [5], [6], [7], [8] |
| $k_{flex}^{hinge}$ | Flexural stiffness of the hinge per unit length | 600 pN·nm | Default value for a single alpha-helix [9], [10], [11] |
| $L$ | Length of the stalk | 5.9 nm | Default value corresponds to the length of the coiled coil fragment in the CG MD model (PDB 3IZ0) [1] |
| $k_{strain}$ | Axial rigidity of 1 nm of the stalk | 1156 pN/nm | Estimated based on the *EI* for a coiled coil [12] |
| $l_0$ | Equilibrium distance between the adjacent stalk sphere centers | 2 nm | Equals the diameter of the spheres, representing a coiled coil |
| $k_{repuls}$ | Stiffness of the steric repulsion between the microtubule and the MAP | 500 pN/nm | This work |
| $F$ | External force | varied:<br>±8, ±16, ±25 pN | This work |
| $T$ | Temperature | 310 K | This work |
| $\eta$ | Dynamic viscosity of the medium | 4.8 mPa·s | This work |
| $\Delta t$ | Time step | 0.02 ns | This work |

### Supplementary figures

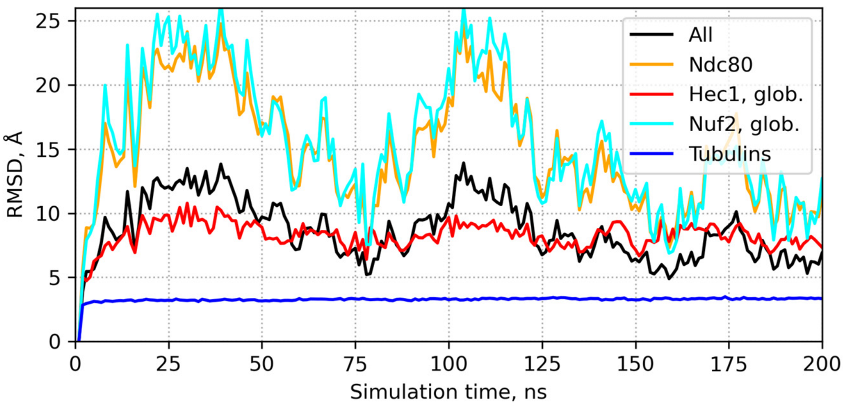


**Figure S1: RMSD of the NDC80 complex in the equilibration simulation.** RMSD is shown as a function of the simulation time for the whole system (black), tubulins (red), the NDC80 complex (orange), and its two fragments: Hec1 (red) and Nuf2 (cyan). Zero load was applied to NDC80 during equilibration. The microtubule remains stable throughout the simulation due to the harmonic positional restraints applied to the inner regions of tubulins. The NDC80 complex undergoes oscillations in RMSD between 7.3 and 24.9 Å from the initial structure with the mean value of 15.4 Å. These undulations are more prominent for the microtubule-distant Nuf2 protein and they mainly occur as a whole-body movement around the Hec1 pivot, which remains more static with the average RMSD of 8.2 Å. Noteworthy, we did not observe any contacts between the globular domain of Nuf2 and the microtubule throughout the simulation.

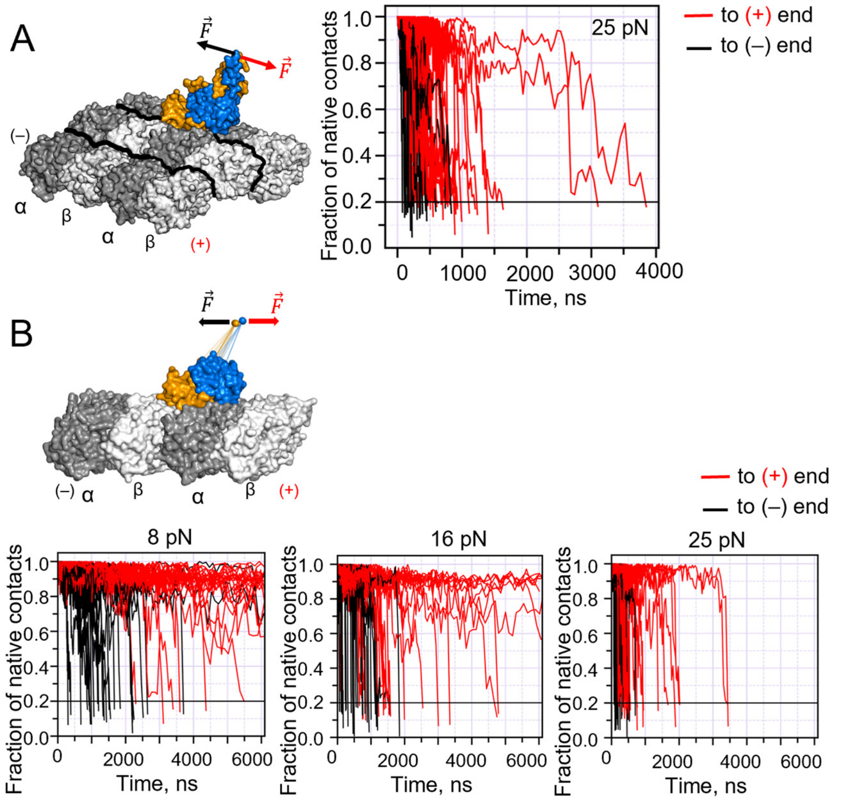


**Figure S2: Extended data on simulation of the unbinding of the NDC80 complex (Hec1/Nuf2 subunits) from the microtubule. (A)** Kinetics of rupture of the native contacts between the Hec1 subunit and the microtubule under plus- and minus-end-directed load of 25 pN, applied to the point of origin of the stalk. Each curve corresponds to an individual simulation run. The rupture time here and below is defined as the time when the number of residual native contacts hits the threshold of 20%. **(B)** Analogous data for the case when the forces of 8 pN, 16 pN, or 25 pN are applied to the tip of the ~6 nm-long virtual stalk. Each curve corresponds to an individual simulation run.

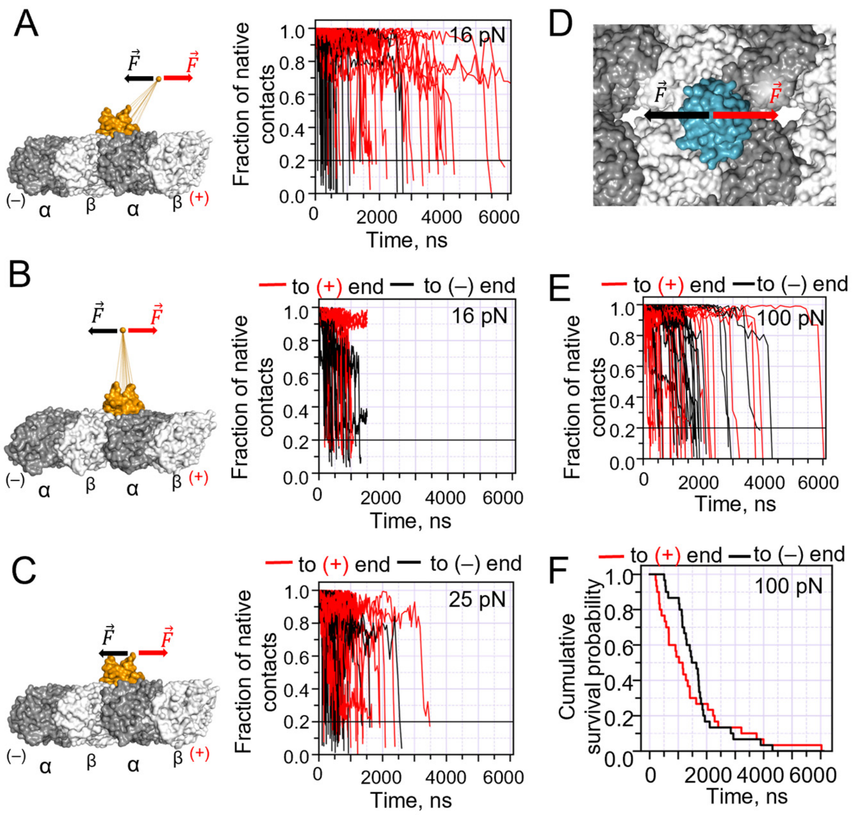


**Figure S3. Extended data on simulation of the unbinding of a single CH domain of Hec1 subunit from the microtubule.** Kinetics of rupture of the native contacts between the Hec1 subunit and the microtubule under plus- and minus-end-directed loads: **(A)** 16 pN, applied to the tip of a ~6-nm-long naturally tilted virtual stalk; **(B)** 16 pN, applied to the tip of orthogonally oriented ~6 nm-long virtual stalk; **(C)** 25 pN, applied to the point of origin of the stalk. **(D)** Schematics of the simulation, in which 100 pN force was applied to the origin of the flexible linker of the CH domain of EB3 protein. **(E)** Kinetics of rupture of the native contacts between the CH domain of EB3 and the microtubule under plus- and minus-end-directed loads. **(F)** Kaplan-Meier plots, showing the cumulative survival probability for residence of the CH domain of EB3 on the microtubule wall in the CG MD simulations under 100 pN load in the plus- and minus-end directions, based on *N* = 30 simulations for each direction of the load.

### Supplementary Video Legends

**Video S1.** CG MD simulations of the NDC80 complex (Hec1/Nuf2 subunits) unbinding from the microtubule when a plus-end-directed force of 25 pN is applied to the tip of the stalk.

**Video S2.** CG MD simulations of the NDC80 complex (Hec1/Nuf2 subunits) unbinding from the microtubule when a minus-end-directed force of 25 pN is applied to the tip of the stalk.

**Video S3.** BD simulations of a MAP, parameterized to reproduce the dimensions, characteristics, and behavior of the NDC80 complex in the CG MD simulations. The applied force is 16 pN in magnitude.

**Video S4.** BD simulations of a MAP, parameterized to have a flexible linker, like in the case of EB proteins. The applied force is 16 pN in magnitude.

**Video S5.** BD simulations of a MAP, parameterized to have the stalk dimensions mimicking a full-length NDC80 complex. The applied force is 25 pN in magnitude.

**Video S6.** BD simulations of a MAP, parameterized to have an orthogonally oriented stalk and an asymmetric distribution of the depths of binding potentials at the microtubule-binding interface (12 $k_BT$, 5 $k_BT$, 5 $k_BT$ from left to right in this video). The magnitude of the applied force is 16 pN.
